# Experimental Bacterial Co-infection in Nile Tilapia Shows High Pathogenicity from Lake Kariba Isolates

**DOI:** 10.1101/2025.03.30.646194

**Authors:** Frederick Chitonga Zulu, Kunda Ndashe, Katendi Changula, Mazuba Siamujompa, Chanda Chitala, Mwansa M. Songe, Ladslav Moonga, Stephen R. Reichley, Bernard Mudenda Hang’ombe

## Abstract

**Objective:** This study aimed to determine the pathogenicity of bacteria (*Aeromonas* spp., *Lactococcus garvieae, Acinetobacter* spp., and *Klebsiella* spp.) isolated from Nile tilapia (*Oreochromis niloticus*) in small-scale aquaculture establishments on Lake Kariba through experimental infections.

**Materials and methods:** Healthy fish (50g ± 5g) were distributed among six transparent fish tanks labeled A to F. Fish in tanks A, B, C, and D were exposed to *Aeromonas* spp., *Lactococcus garvieae, Acinetobacter* spp., and *Klebsiella* spp., respectively, through intraperitoneal inoculation. Tank E received a co-infection of all isolates, while tank F served as the control group with an injection of normal saline. Clinical signs, mortalities and post-mortem lesions were recorded in all experimental groups, with histopathological examinations performed on liver, kidney, and spleen tissues.

**Results:** The findings indicate that *Acinetobacter* spp. and *Klebsiella* spp. exhibit low pathogenicity, evidenced by few clinical signs such as lethargy, pale skin, and fin erosion. The study further highlighted *Aeromonas* spp. and *Lactococcus garvieae* as the bacterial isolates causing significant clinical symptoms in Nile tilapia compared to *Acinetobacter* spp. and *Klebsiella* spp. following experimental infection.

**Conclusion:** Co-infection with all bacterial pathogens demonstrated very high pathogenicity, with 100% mortality reported within seven days post-infection. The study proposes the development of a polyvalent vaccine to control fish disease outbreaks in small-scale aquaculture operations on Lake Kariba, thereby sustaining aquaculture production in Zambia.

## 1. Introduction

Globally, aquaculture production growth is driven by both large commercial operations and small-scale producers (1). Over the past two decades, global aquaculture production has increased at an annual rate of 6.7%, making it the fastest-growing sector in animal production (1). However, this growth declined to 3.5% for the period 2016 to 2021 (2). In some parts of the world the significant factor contributing to this decline is disease outbreaks in production facilities (3,4).

Globally, the production of Nile tilapia (*Oreochromis niloticus*) is particularly affected by disease outbreaks, predominantly bacterial in nature (5,6). Reported diseases in farmed Nile tilapia include francisellosis (*Francisella noatunensis* subsp. orientalis), lactococcosis (*Lactococcus garvieae*), streptococcosis (*Streptococcus agalactiae, S. iniae*, and *S. dysagalactiae*), edwardsiellosis (*Edwardsiella tarda* and *E. ictaluri*), and motile hemorrhagic septicemia (*Aeromonas hydrophila, A. sobria*, and *A. veronii*) (7–11). These bacterial infections often emerge following exposure to stressors such as overcrowding, poor water quality, and suboptimal nutrition, which suppress fish immunity and increase susceptibility to environmental bacteria. Consequently, it is common to isolate multiple bacterial pathogens during an outbreak in farmed tilapia (12).

In Zambia, aquaculture production has recently increased, with significant participation from small-scale producers over the past five years (13). The Government of the Republic of Zambia, through various projects and funding agencies, has promoted financial investment in small-scale fish farming (14). Substantial investment has been directed toward increasing the number of small-scale producers on Lake Kariba, the largest fish production region in Zambia (15). Siamunjopa *et al*. (2023) reported significant mortality losses of fish among small-scale producers on Lake Kariba due to *Aeromonas* spp., *Pseudomonas* spp., *Micrococcus* spp., *Klebsiella* spp., *Lactococcus* spp., *Streptococcus* spp., and *Acinetobacter* spp (16).

Despite the isolation of bacteria from diseased Nile tilapia in small-scale fish farms on Lake Kariba, the pathogenicity of these isolates remains unknown. It is well documented that immunosuppressed fish, due to poor husbandry practices, are susceptible to multiple bacterial pathogens (12,17,18). Therefore, this study aimed to determine the pathogenicity of the bacterial isolates from diseased Nile tilapia from small-scale producers to elucidate the role of these bacterial pathogens in the disease process.

## 2. Materials and Methods

This study was undertaken in strict accordance with the recommendations in the Guide for the Care and Use of Laboratory Animals of the National Health Research Ethics Committee of Zambia. The protocol of this study was approved by the Excellence in Research Ethics and Science (ERES) Converge, (Protocol Number: 2019/AUG/024). All efforts were made to minimize suffering and stress of the fish, both during handling and sampling. As infection was one of the humane endpoints, subjects were withdrawn from the experiment (euthanized and sampled) after clinical signs appeared. The decision criteria to euthanize animals included two or more clinical signs (e.g. poor body condition, severe skin erosion and hemorrhage, loss of balance, extensive abdominal swelling, scale protrusion, and exophthalmia). Where signs of disease or abnormal behaviour were observed, the fish were euthanised by stunning with a blow to the head followed by dislocation of the cervical vertebra.

### 2.1. Animals and environmental conditions

Healthy *Oreochromis niloticus* (Nile tilapia) juveniles with average 40g ± 12.5g body weight were obtained from Palabana Fisheries, a commercial farm without history of disease outbreak, located in Chirundu district, South-East of Zambia. The fish population in this farm had been observed for clinical signs and sampled for evidence of disease over a period of 6 months. The fish were maintained in 500-L tanks for at least 2 weeks before conducting the experiments. The animals were maintained on commercial feed which was supplied twice a day to satiety. Feeding was withdrawn 24 h before experiments, and the animals were anesthetized in 10 mg L of tricaine methanesulfonate (MS222) before bacterial administration. Water parameters such as dissolved oxygen (DO), temperature, pH, ammonia, total dissolved solids and electrical conductivity were continuously measured.

### 2.2. Bacterial Pathogens

Bacterial pathogens used in this study were *Aeromonas spp*., *Lactococcus garvieae*., *Acinetobacter* spp., and *Klebsiella* spp. e previously isolated from a natural outbreak of disease in Nile tilapia in small cage cultured fish on Lake Kariba, Siavonga, Zambia (16). The bacteria were cultured in brain–heart infusion (BHI) agar enriched with 5% defibrinated ovine blood and incubated at 28 °C for 24 h. Individual isolates of each bacteria were grown in BHI broth, and stocks were made with 15% glycerol and kept at −80 °C until use.

### 2.3. Determination of Lethal Dose (LD_50_) of the bacteria

An experimental challenge study was conducted to find out the lethal dose - 50% end point (LD50) in Nile tilapia. The bacteria were subcultured in 10 ml of Brain Heart Infusion (BHI) broth and were incubated at 37 ºC for 24h. After 24 h incubation, the stock solutions were prepared by dissolving colonies of each bacterial isolate in saline solution to match the turbidity of four McFarland standards (Riga, Latvia) [Table 1].

**Table 1:** McFarland standards.

| S/N | McFarland Standard | Approximate Cell Count Density (x10<sup>8</sup> cells) |
| --- | --- | --- |
| 1 | 0.5 | $1.5 \times 10^8$ |
| 2 | 1 | $3.0 \times 10^8$ |
| 3 | 2 | $6.0 \times 10^8$ |
| 4 | 3 | $9.0 \times 10^8$ |
| 5 | 4 | $12.0 \times 10^8$ |

The test was conducted with batches of 10 fish per dose by intraperitoneal injection in Nile tilapia with 24 hrs of bacterial suspension. The fishes were injected with 0.1 ml of bacterial suspension intraperitoneally with final concentrations of 1.5 × 10^8^, 3.0 × 10^8^, 6.0 × 10^8^, 9.0 × 10^8^ and 12.0 × 10^8^ CFU per ml, respectively. The control fishes were injected with 0.2 ml of normal saline. The challenge study was carried out in triplicates. A total number of 20 fishes were kept in each tank (40L). Fish mortality was recorded in every 24 h interval for 10 days. LD_50_ was calculated using the Reed and Muench (1938) method.

### 2.4. The Challenge Study

The study had six treatment groups: *Aeromonas spp*., *Lactococcus garvieae, Acinetobacter* spp., *Klebsiella* spp, co-infection of all bacteria, and negative control. Twenty(20) fish per treatment group were inoculated intraperitoneally with 100 μl bacterial suspension at the determined lethal doses according to the treatment groups [Table 2].

**Table 2:** Treatment groups for the trial experiments.

| Experiment Group | Isolate Name | LD<sub>50</sub> (CFU/ Fish) | Volume |
| --- | --- | --- | --- |
| Group A | *Klebsiella* spp | 1.05 x 10<sup>6</sup> | 100 µl |
| Group B | *Acinetobacter* spp | 1.80 x 10<sup>6</sup> | 100 µl |
| Group C | *Aeromonas* spp | 8.5 x 10<sup>5</sup> | 100 µl |
| Group D | *Lactococcus garvieae* | 9.6 x 10<sup>5</sup> | 100 µl |
| Group E | *Klebsiella* spp | 1.05 x 10<sup>6</sup> | 100 µl |
|  | *Acinetobacter* spp | 1.80 x 10<sup>6</sup> |  |
|  | *Aeromonas* spp | 8.5 x 10<sup>5</sup> |  |
|  | *Lactococcus garvieae* | 9.6 x 10<sup>5</sup> |  |
| Group F | Control (Normal Saline) | Normal Saline | 100 µl |

Clinical signs after bacterial inoculation were monitored daily for 21 days and the gross pathology and mortality rate were recorded daily. Necropsy, bacteriological, and histological examination was conducted on moribund fish r. At 21 days post-infection, all surviving fish were euthanized using an overdose of eugenol solution.

### 2.5. Histopathological examination of challenged fish

Tissue samples (Anterior kidney, liver, and spleen) from the control and infected groups (three per group) were collected for histological study. These samples were preserved in 10% (vol/vol) neutral buffered formalin. After 24 h, the formalin was replaced with a fresh 10% formalin solution. Tissue sections were embedded with paraffin and stained with hematoxylin and eosin (H&E) using standard histological procedures.

### 2.6. Bacteriological examination of challenged fish

To satisfy Koch’s postulates the bacteria was reisolated and identified from the moribund fishes. The brain, liver, and kidney were sampled using sterile inoculation loops and streaked on MacConkey agar (HiMedia, India), nutrient agar (HiMedia, India), and blood agar (HiMedia, India) using a strictly aseptic technique. The inoculated Petri dishes were stored at 28 °C for 24 h.

Pure cultures were obtained by carrying out subculturing procedures and subjecting them to a second round of incubation at room temperature for another 48 hours, ensuring the isolation of uncontaminated bacterial strains. The isolates were identified by determining colony morphology and afterward the isolates were grouped accordingly. Two to three representative isolates from each group were subjected to Gram-staining. Conventional biochemical tests to characterize the bacteria was done according Siamunjompa *et al*., (2023) (16).

### 2.7. Data Analysis

Data were entered into an Excel spreadsheet (Microsoft Excel 2010 version, Redmond, WA, USA) and then exported to DATA Tab™ (Styria, Austria), a Web-App for statistical data analysis, where descriptive statistics (frequencies and proportions) were computed and presented using tables for categorical parameters (19). The summary tables and graphs were prepared in accordance with the objective of the study.

## 3. Results

### 3.1. Observation of clinical signs

The clinical signs observed in the days following infection indicated that in fish infected with *Klebsiella* and *Acinetobacter*, the initial signs appeared 2 days post-infection [Figure 1]. *Klebsiella*-infected fish exhibited lethargy, skin ulcerations, and mortality, while *Acinetobacter*-infected fish showed pale skin, loss of appetite, and mortality [Figure 1]. In contrast, the first clinical signs in fish infected with *Aeromonas* spp and *Lactococcus garvieae* were recorded 1 day post-infection [Figure 1]. *Aeromonas*-infected fish displayed hemorrhages on the body and trunk, skin ulcerations, ascites, and loss of appetite and mortality? [Figure 1]. Fish infected with *Lactococcus garvieae* exhibited erratic swimming, fin erosion, lethargy, corneal opacity, skin ulcerations, and mortality [Figure 1]. For the co-infected group, significant clinical signs included loss of appetite, body hemorrhages, skin ulcerations, fin hemorrhages, and high mortality [Figure 1].

**Figure 1:**
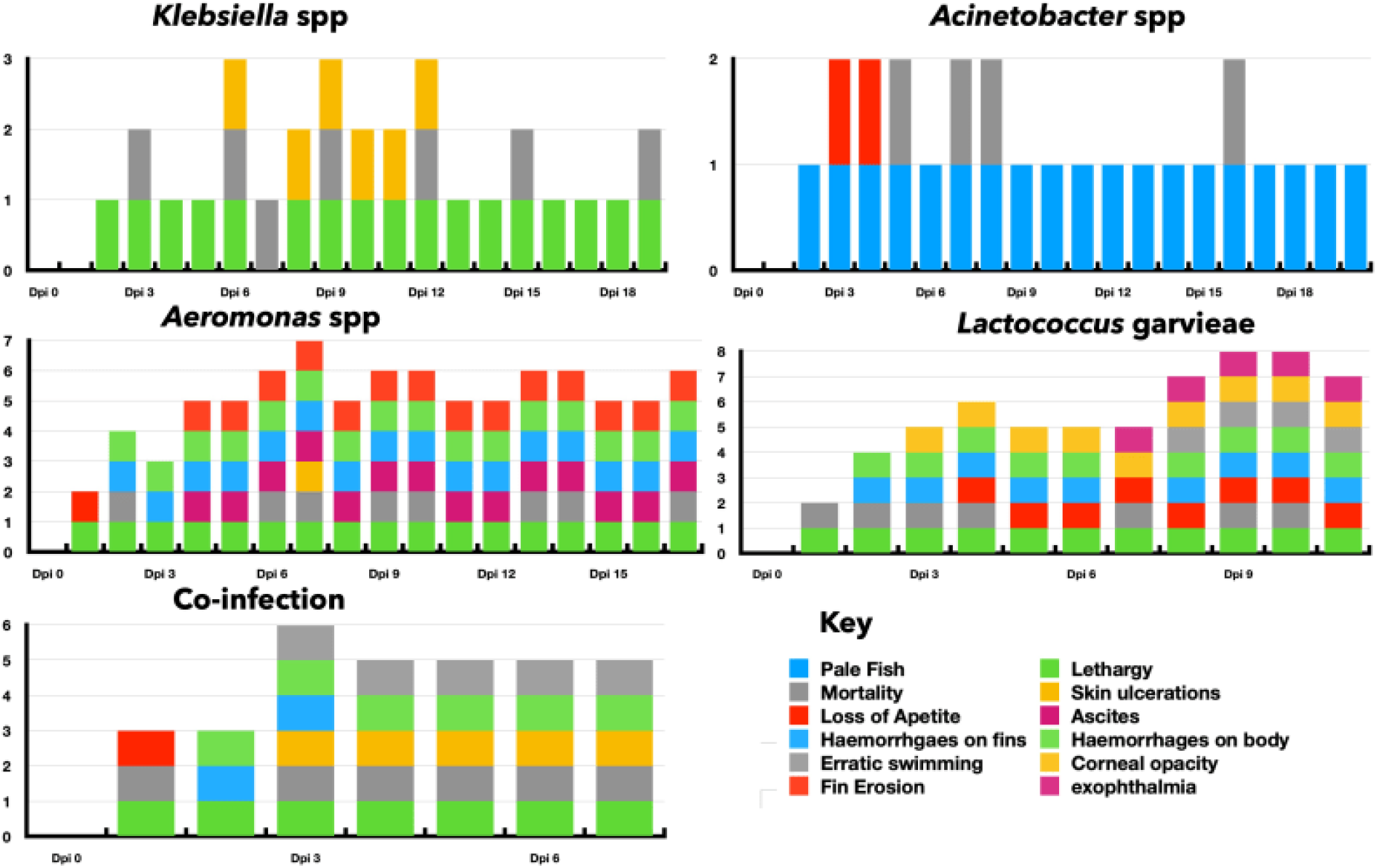
Clinical signs observed on fish infected with the various bacteria in infection experiments

### 3.2. External pathological lesions

The external pathological lesions observed in the experimental groups are detailed in Figure 2. In the *Aeromonas* group, there were slight erosions, minor skin ulcerations, and hemorrhaging on the operculum and eyes (Figure 2A). The *Klebsiella* group exhibited hemorrhages on the trunk and fins (Figure 2B). Fish in the *Acinetobacter* group appeared pale with slight hemorrhages on the operculum (Figure 2C). The *Lactococcus garvieae* group showed corneal opacity, missing scales, fin erosion, and hemorrhages on the operculum (Figure 2D). The co-infection group had skin ulcerations, fin erosions, hemorrhages on the operculum, and wounds around the mouth (Figure 2E).

**Figure 2:**
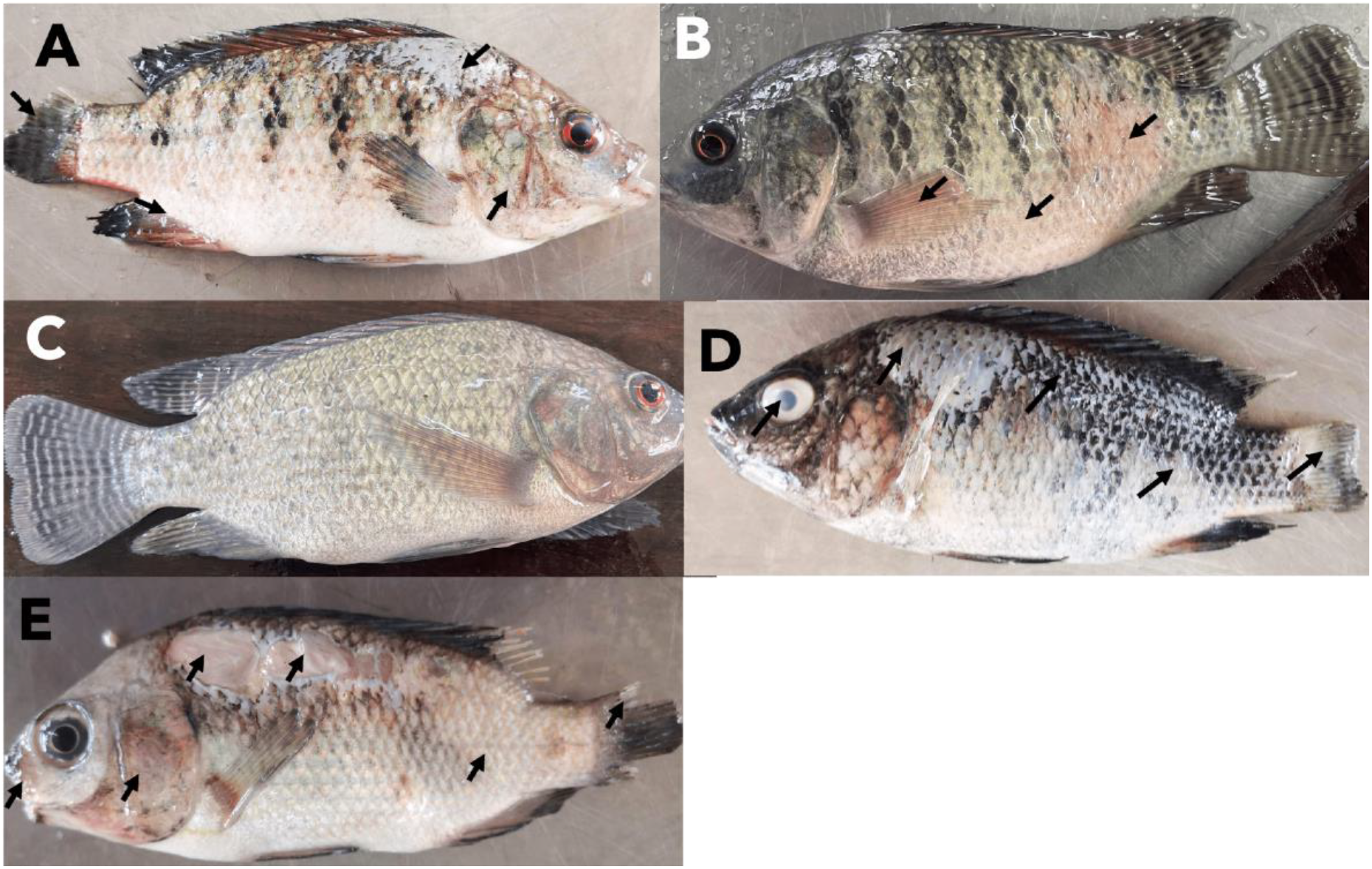
The external pathological lesions observed in the experimental groups were as follows: In the *Aeromonas* group (A), there were slight erosions, skin ulcerations, and hemorrhaging on the operculum and eyes. The *Klebsiella* group (B) exhibited minor skin ulceration, and hemorrhages in the eyes, on the trunk, and fins. Fish in the *Acinetobacter* group (C) appeared pale with slight hemorrhages in the eyes and on the operculum. The *Lactococcus garvieae* group (D) showed corneal opacity, missing scales, fin erosion, and hemorrhages on the operculum. The co-infection group (E) had skin ulcerations, fin erosions, hemorrhages on the operculum, and wounds around the mouth.

### 3.3. Histopathological assessments

The liver of fish infected with *Klebsiella* spp exhibited congestion and hemorrhages (Figure 3A, B, C), parenchymal degeneration (Figure 3B), and lymphocyte infiltration (Figure 3D). In fish infected with *Acinetobacter* spp, the liver showed congestion (Figure 4A), parenchymal degeneration (Figure 4B), and hepatocyte vacuolation (Figure 4B), while the spleen revealed eosinophil infiltration (Figure 4C) and melanomacrophage centers (Figure 4C). Fish infected with *Aeromonas* spp presented main lesions of parenchymal degeneration (Figure 5A), hepatocyte vacuolation, and lymphocyte infiltration (Figure 5A). In fish infected with *Lactococcus garvieae*, significant liver lesions included congestion (Figure 6A), lymphocyte infiltration (Figure 6B, D), and parenchymal degeneration (Figure 6D), with inflammatory cell infiltration observed in the epidermis (Figure 6C, F). Co-infected fish displayed significant lesions across the liver, kidney, skin, and spleen. In the kidney, there was excessive lymphocyte infiltration (Figure 7A, B), parenchymal necrosis and degeneration (Figure 7B). The liver showed lymphocyte infiltration (Figure 7C, D), parenchymal degeneration (Figure 7D), and hepatocyte vacuolation (Figure 7D). In the skin, inflammatory cell infiltration was noted, while the spleen exhibited eosinophil infiltration (Figure 7F) and parenchymal degeneration (Figure 7E).

**Figure 3:**
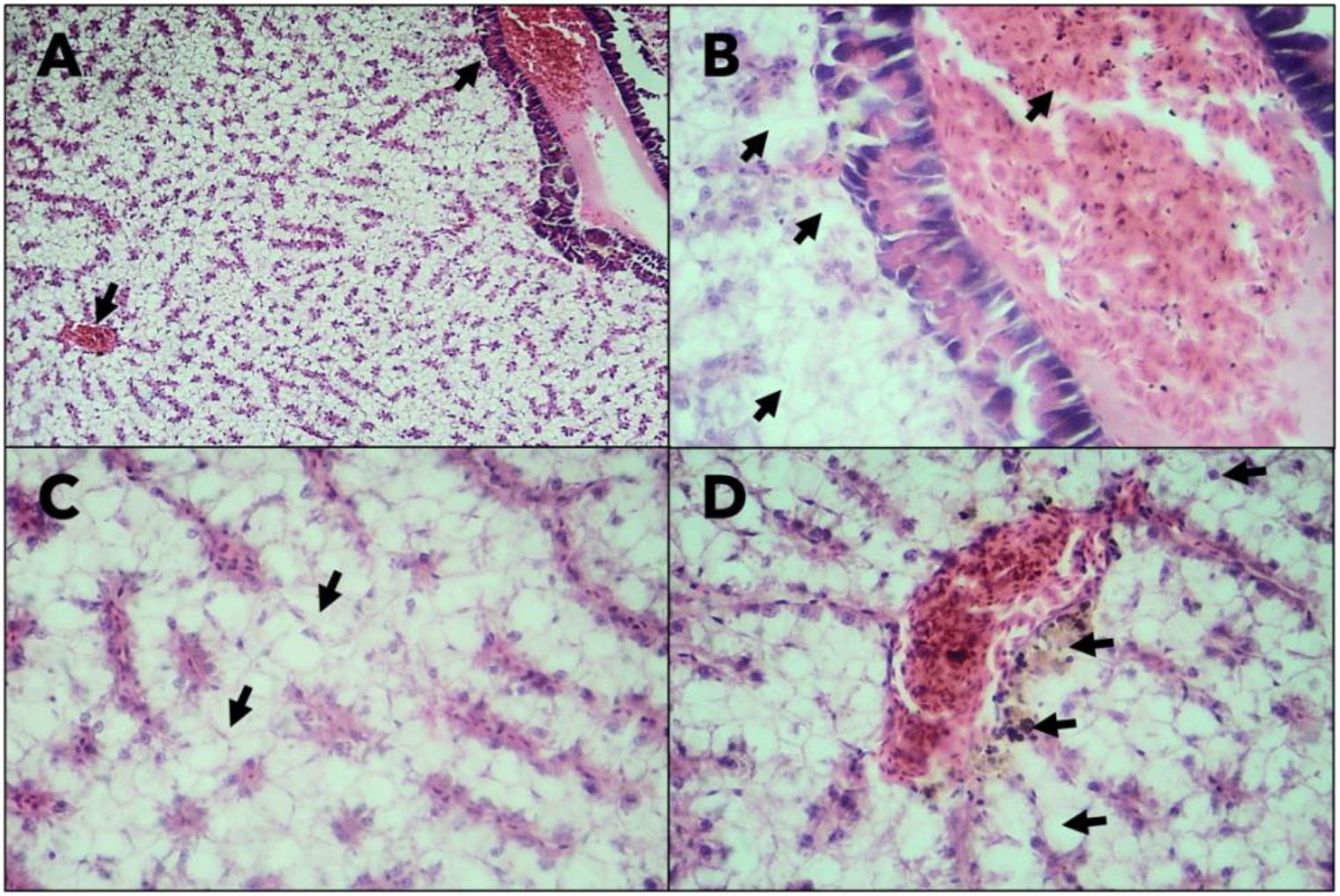
Liver of fish infected with *Klebsiella* spp showing congestion and haemorrhages (A, B and C), degeneration of parenchyma (B) and lymphocyte infiltration (D).

**Figure 4:**
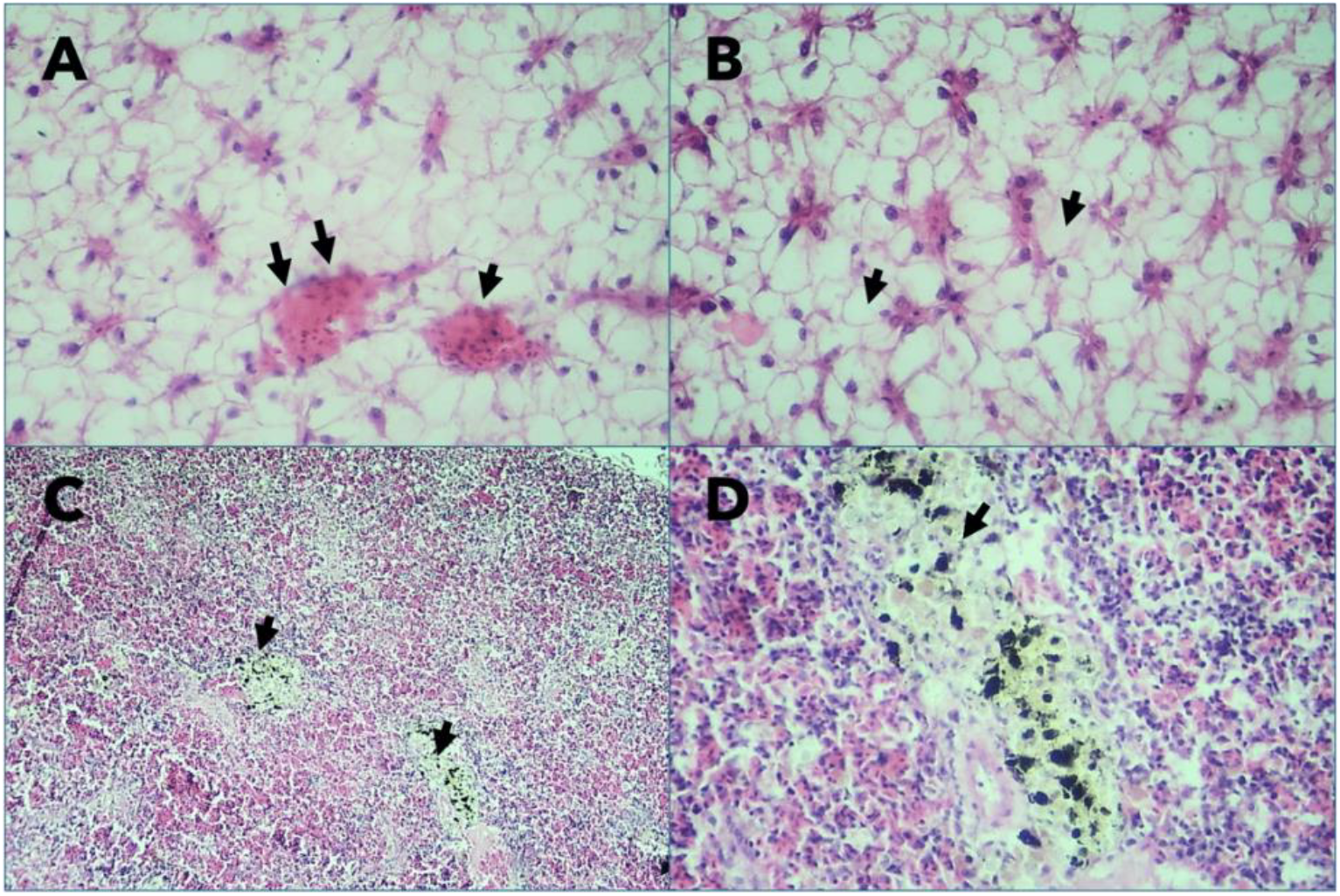
In the liver of Fish infected with *Acinetobacter* spp showed congestion (A) and degeneration of parenchyma (B), and vacuolation of hepatocytes (B). In the spleen, eosinophil infiltration (C) and centres of melanomacrophages were seen.

**Figure 5:**
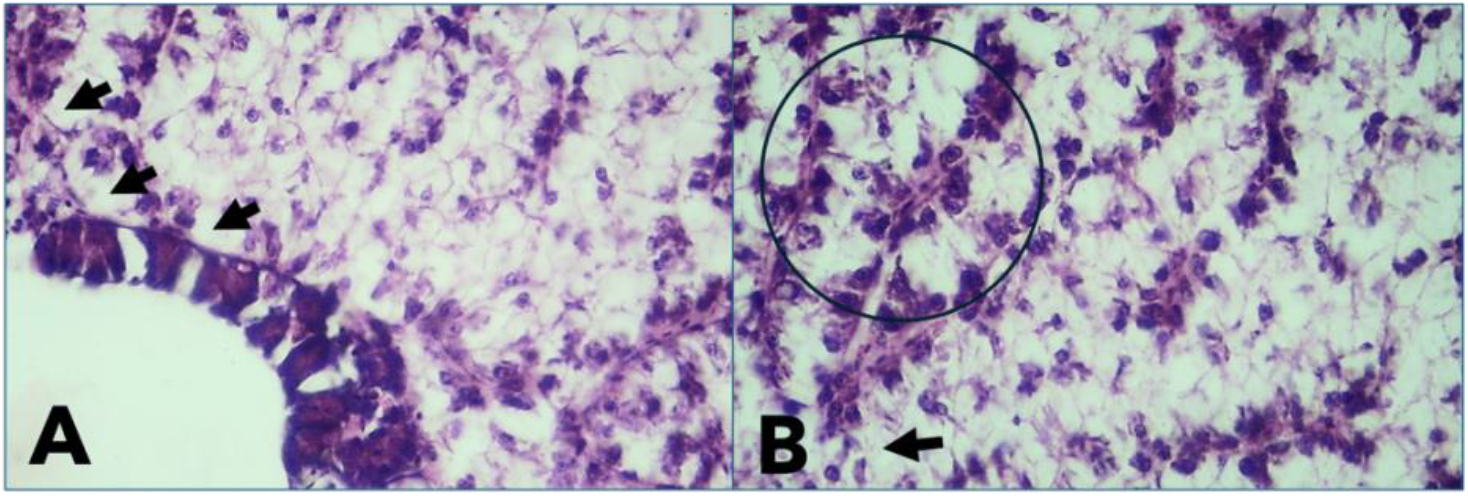
in the fish infected with *Aeromonas* spp, the main lesions were degeneration of the parenchyma (A), vacuolation of hepatocytes, and lymphocyte infiltration.

**Figure 6:**
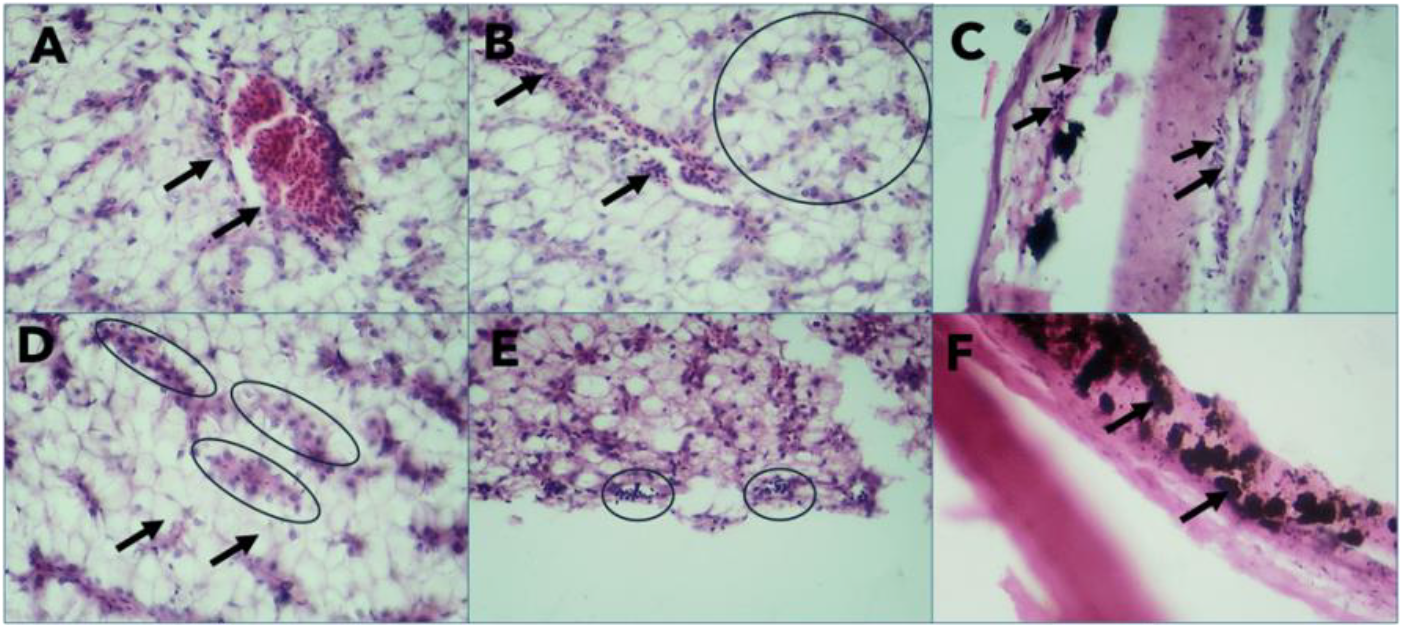
In the fish infected with Lactococcus garvieae in the liver the significant lesions were congestion (A), infiltrations of lymphocytes (B and D), and degeneration of parenchyma (D). in the epidermis, the significant lesions were infiltration of inflammatory cells (C and F).

**Figure 7:**
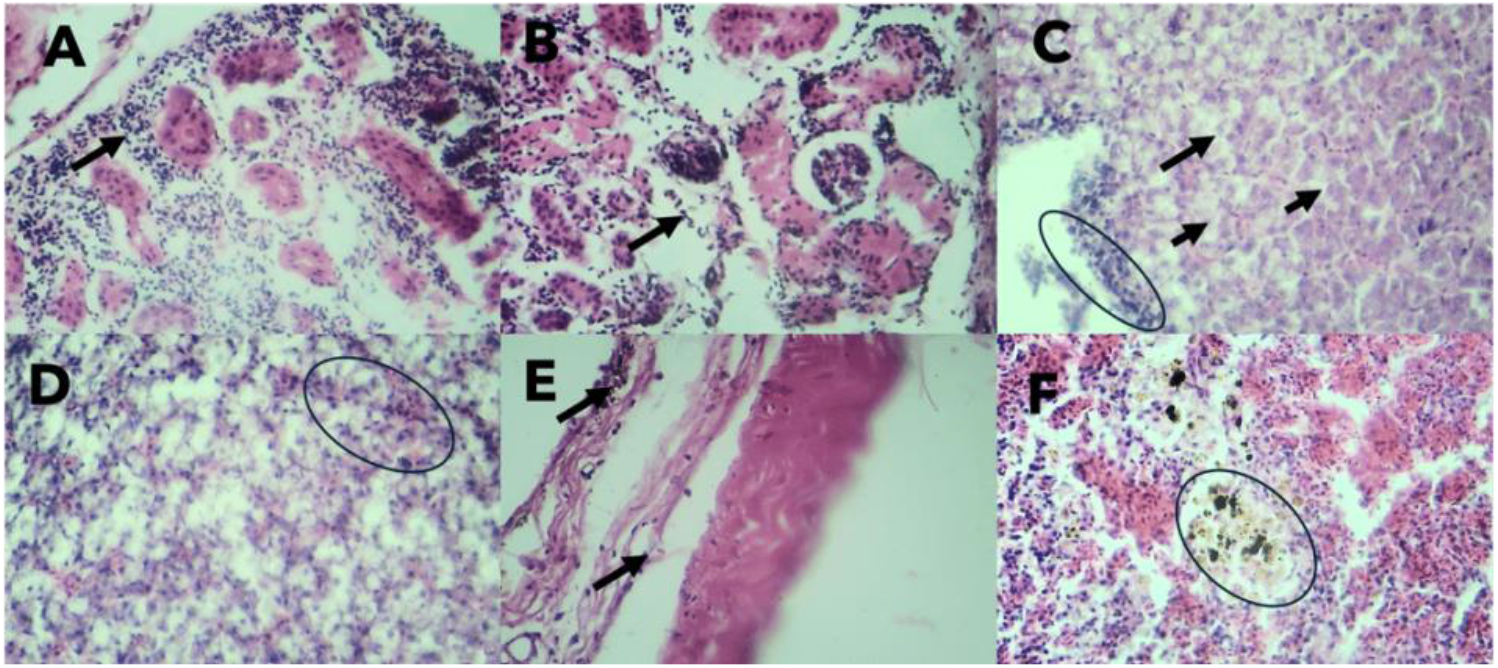
the co-infected fish has significant lesions seen in liver, kidney, skin and spleen. In the kidney excessive infiltration of lymphocytes (A and B), and necrosis and degeneration of parenchyma (B). In the liver, infiltration of lymphocytes (C and D), degeneration of parenchyma (D), and vacuolation of hepatocytes (D). In the skin the significant lesion was infiltration of inflammatory cells. In the spleen, eosinophil infiltration (F) and degeneration of parenchyma (E) were observed.

### 3.4. Mortality trends in the different experimental groups

Comparative analysis of cumulative mortality revealed the most rapid demise in the co-infection group, with 100% (20/20) mortality by 7 days post-infection (dpi) [Figure 8]. In the *Lactococcus garvieae*-infected group, initial mortality was observed at 2 dpi, and all succumbed by day 12 dpi [Figure 8]. *Aeromonas* spp. infection also exhibited initial mortality at 2 dpi, with complete mortality by day 16 dpi [Figure 8]. Conversely, the *Klebsiella* and *Acinetobacter* groups displayed the lowest mortality rates with 7 and 5 mortalities out of twenty recorded respectively at the study’s conclusion [Figure 8].

**Figure 8:**
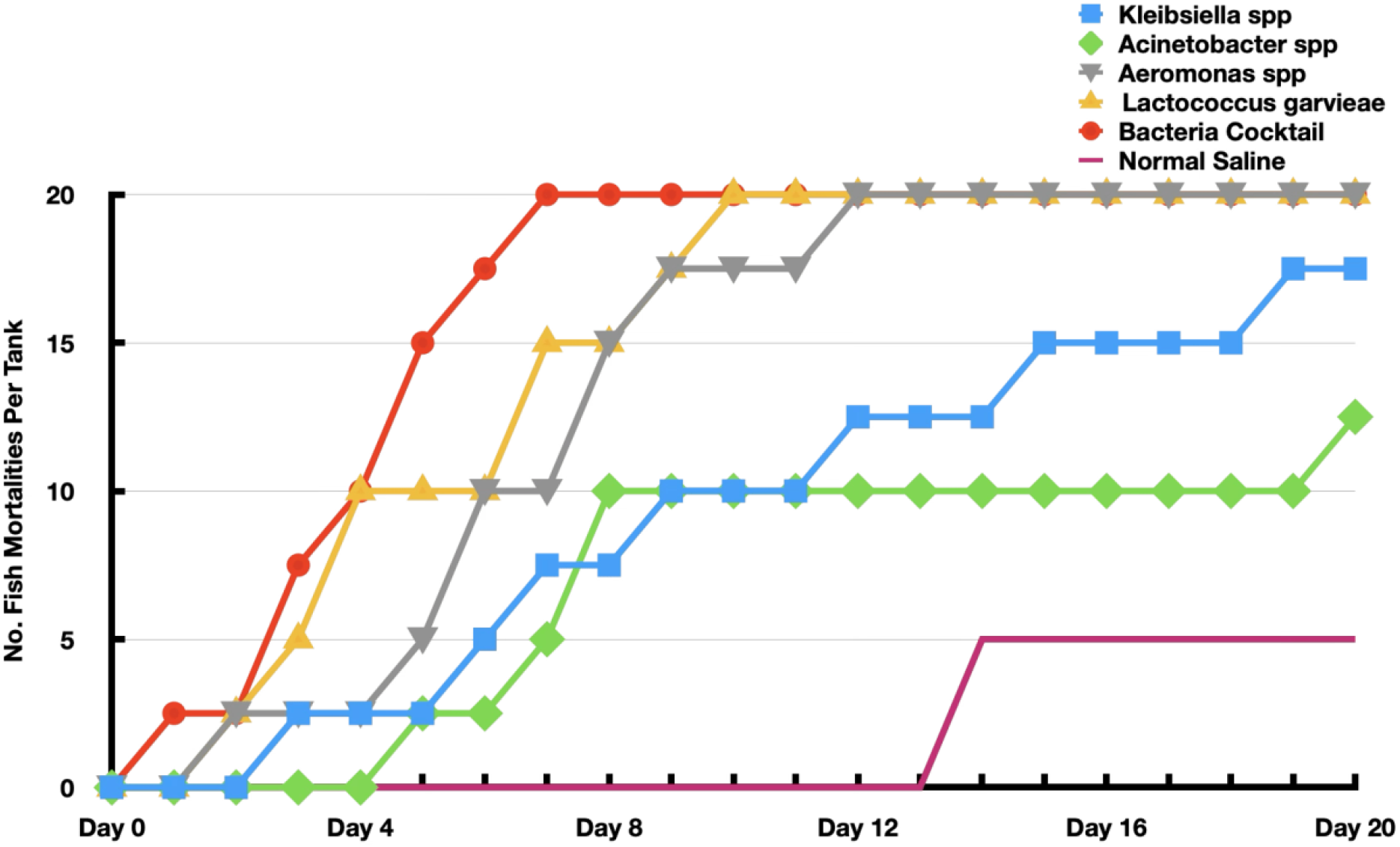
Number of mortalities of fish challenged with indicated bacteria isolated in this study.

### 3.5. Bacteriological examination of challenged fish

Bacterial isolation from the experimental fish revealed that in groups A and B, the experimental bacteria *Klebsiella* spp and *Acinetobacter* spp were isolated exclusively from the liver [Table 3]. In groups C and D, the experimental bacteria *Aeromonas* spp and *Lactococcus* garvieae were isolated from the spleen, liver, and kidney [Table 3]. Additionally, *Lactococcus garvieae* was isolated from brain tissue. In the co-infection group, *Aeromonas* spp and *Lactococcus* garvieae were isolated from the spleen, kidney, and brain, while all the bacteria were isolated from the liver [Table 3].

**Table 3:**
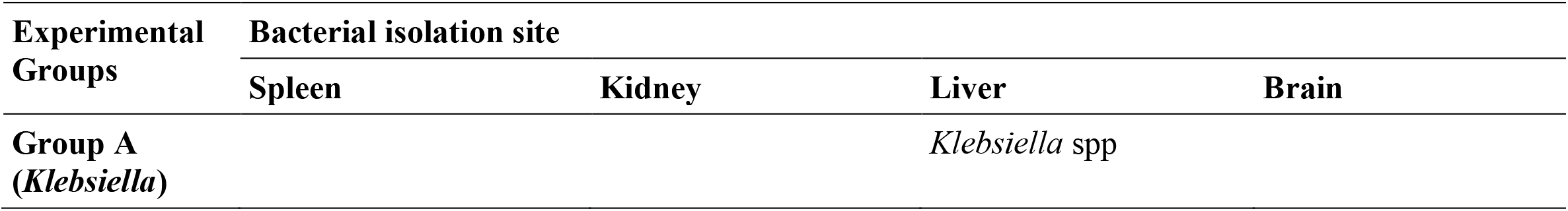
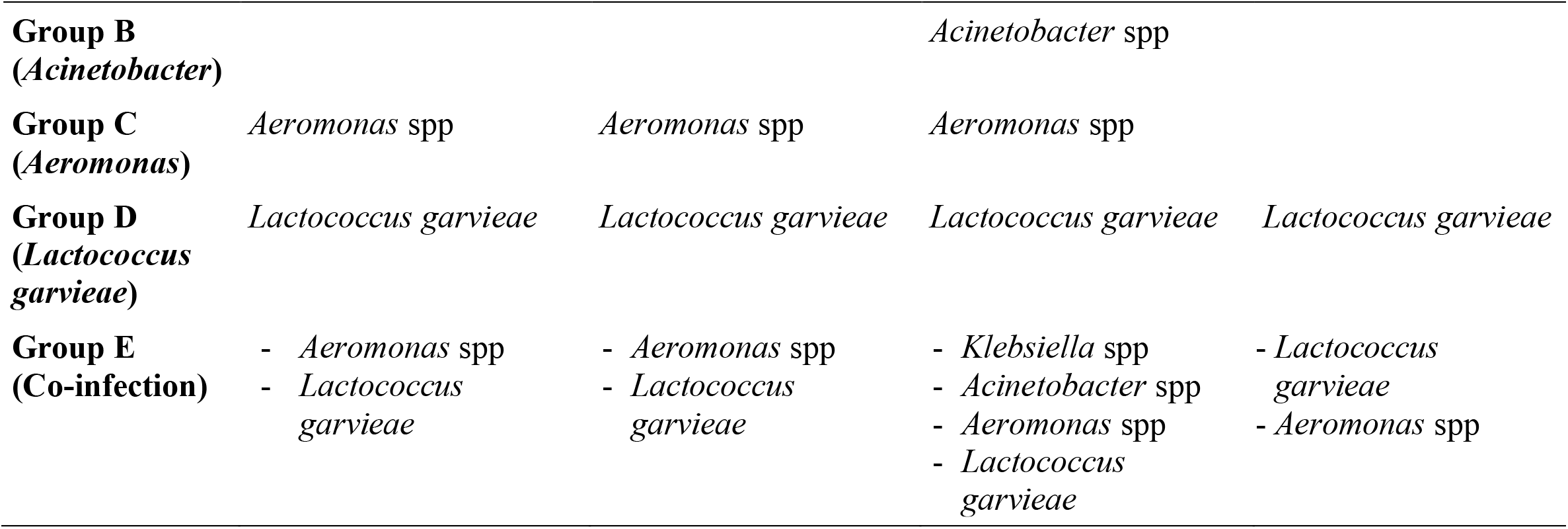
Isolation of bacteria from organs of fish in the infection experiment.

## 4. Discussion

Bacterial pathogens affecting farmed Nile tilapia in both large and small-scale operations on Lake Kariba have been documented (11,16). Fish reared in floating cages are naturally exposed to the microbial communities in the aquatic environment, inevitably leading to co-infections during disease outbreaks. Our study demonstrate that co-infection with four species of bacteria resulted in extremely high pathogenicity among the experimental fish. *Aeromonas* spp and *Lactococcus garvieae*, well-documented fish pathogens (9,10,20), exhibited high pathogenicity, whereas *Klebsiella* spp and *Acinetobacter* spp, which are not well-characterized as fish pathogens, demonstrated reletively low pathogenicity.

Challenge fish receiving *Acinetobacter* spp. exhibited clinical signs of pale skin, loss of appetite, and mortality. These pathogens have not previously been reported to cause significant disease outbreaks in Nile tilapia production globally. However, *Acinetobacter* spp. have been documented to cause disease outbreaks in other fish species, such as rohu (*Labeo rohita*) (21). The clinical signs observed in *Acinetobacter* outbreaks in rohu included ulcerative lesions, hemorrhages on the skin and tail, and fin rot, which differ from the observations in this study where Nile tilapia presented with pale skin and loss of appetite. Histopathological examination in the present study revealed liver lesions, including parenchymal degeneration and hepatocyte vacuolation, and in the spleen, eosinophil infiltration and melanomacrophage centers. These findings were consistent with those reported by Laltlanmawia *et al*. (2023) in diseased rohu (21). The mortality rate at the end of the present experiment, 21 days post-infection, was 50%, in contrast to the infection experiment in rohu, where 100% mortality was recorded within the first 3 days post-infection (21).

In the present study, fish challenged with *Klebsiella* spp. exhibited clinical signs of skin ulcerations, lethargy, and mortality. This pathogen has previously been isolated from diseased tilapia, where significant mortality was recorded (16,22,23). Reported clinical signs in these studies included lethargy, anorexia, subcutaneous hemorrhages, urogenital bleeding, and ascites, some of which were consistent with the present findings (16,22,23). Vaneci-Silva et al. (2022) reported significant histopathological lesions such as multifocal edema and fatty degeneration in the liver, as well as melanomacrophage centers in the spleen and kidney (23). The present study also identified significant histopathological lesions in the liver, including congestion and hemorrhages, parenchymal degeneration, and lymphocyte infiltration. Additionally, the cumulative mortality rate in this study was 70% at 21 days post-challenge, whereas an outbreak in juvenile Nile tilapia at a farm in Brazil resulted in 100% mortality within 4 days (23).

*Aeromonas* spp have been reported globally as significant pathogens in aquaculture, often causing outbreaks secondary to stressors in production systems (9,20,24,25). Clinical signs of *Aeromonas* infections include hemorrhagic patches on the skin, at the base of the pectoral fin, and around the anal opening, as well as scale desquamation, skin ulcerations and fin erosions (20,24). These signs were consistent with those observed in the present study. Histopathological examination of liver tissues in this study revealed parenchymal degeneration, hepatocyte vacuolation, and lymphocyte infiltration, consistent with findings by Dong et al. (2017), who additionally reported hemosiderin accumulation around vessels and hepatocytes, and hyperemia and hemorrhage in the spleen (9). The cumulative mortality rate in the present study was 100% at 16 days post-challenge, contrasting with the 70% cumulative mortality rate at 21 days post-challenge reported by Dong *et al*., (2017).

Since the initial isolation of *Lactococcus garvieae* from Nile tilapia in Zambia in 2018, this pathogen has continued to be reported as a cause significant losses due to disease outbreaks among both small and large commercial producers (11,16,22). The main clinical signs observed in the present study, such as erratic swimming, corneal opacity, fin erosion, and loss of scales, are consistent with those reported in outbreaks among farmed tilapia in Zambia (11,16). Histopathological lesions recorded in this study included congestion, lymphocyte infiltration, parenchymal degeneration, and inflammatory cell infiltration in the skin epidermis. These lesions are similar to those reported in cage-cultured cobia (*Rachycentron canadum*) (26). The present study recorded a cumulative mortality rate of 100% by 12 days post-infection which highlights the pathogenic nature of the bacterial causing significant mortality in many tilapia production farms, both small and large-scale, in Zambia (11,16).

Previous experimental studies on co-infection in Nile tilapia primarily focused on bacteria, viruses, or parasites (9,27,28). This study demonstrated the co-infection of four bacterial pathogens, previously isolated from farmed Nile tilapia on Lake Kariba among small-scale producers (16). To the best of our knowledge, this is the first study to elucidate the co-infection of four bacterial pathogens in Nile tilapia. Clinical signs were observed within a day post-infection, earlier than for any of the single pathogens included in the study. These signs included skin ulcerations, mouth lesions, fin erosions, corneal opacity, and hemorrhages, with the first three lesions being more severe in the co-infection group than in the mono-infection experiments. Dong et al. (2017) reported heavy mortalities in Nile tilapia infected with *Aeromonas jandaei* and *Aeromonas veronii* (9). In the present study, co-infection with four bacterial pathogens resulted in very high mortality, with cumulative mortality reaching 100% by the seventh day post-infection. Histopathological examination further demonstrated the pathogenicity of the co-infection, showing a combination of lesions seen in the mono-infections. These lesions included excessive lymphocyte infiltration, necrosis, and parenchymal degeneration in the kidney; lymphocyte infiltration, parenchymal degeneration, and hepatocyte vacuolation in the liver; inflammatory cell infiltration in the skin; and eosinophil infiltration and parenchymal degeneration in the spleen. The results of this study indicate that the presence of *Klebsiella* spp., *Acinetobacter* spp., *Aeromonas* spp., and *Lactococcus garvieae* in the cage environment, coupled with stressors, will lead to severe mortalities in farmed fish.

This study demonstrated that Nile tilapia infected with *Klebsiella* spp. and *Acinetobacter* spp. exhibited low pathogenicity. Furthermore, we reported for the first time the experimental co-infection of Nile tilapia with four bacterial pathogens (*Klebsiella* spp., *Acinetobacter* spp., *Aeromonas* spp., and *Lactococcus garvieae*). The high pathogenicity observed in these bacterial co-infections suggests a potential threat to the aquaculture industry in Zambia. Therefore, future research should focus on developing strategies to control these bacterial co-infections, including the development of polyvalent vaccines.

## 5. Acknowledgements

The authors would like to thank the technical and administrative staff of the University of Zambia, School of Veterinary Medicine, Bacteriology and Pathology Units who supported and faciliated the study.

## 6. Statements and Declarations

### Authors’ contributions

All authors contributed to the study conception and design. Material preparation, data collection and analysis were performed by Kunda Ndashe, Frederick Chitongo Zulu, Ladslav Moonga and Katendi Changula. The first draft of the manuscript was written by Kunda Ndashe, and Frederick Chitongo Zulu and all authors commented on previous versions of the manuscript. All authors read and approved the final manuscript

### Data Availability

The data that support the findings of this study are available from the corresponding author, Kunda Ndashe, upon reasonable request.

### Ethics approval

This study was undertaken in strict accordance with the recommendations in the Guide for the Care and Use of Laboratory Animals of the National Health Research Ethics Committee of Zambia. The protocol of this study was approved by the Excellence in Research Ethics and Science (ERES) Converge, (Protocol Number: 2019/AUG/024).

### Statement of Animal Rights

All procedures involving animals were conducted in compliance with ethical guidelines and approved by the relevant institutional ethics committee. The welfare of the animals was ensured throughout the study, adhering to the principles of humane treatment, minimizing distress, and following the 3Rs (Replacement, Reduction, and Refinement) wherever applicable.

## References

1. FAO. The State of World Fisheries and Aquaculture 2022: Towards Blue Transformation [Internet]. Rome, Italy: FAO; 2022 [cited 2022 Dec 19]. 266 p. (The State of World Fisheries and Aquaculture (SOFIA)). Available from: https://www.fao.org/documents/card/en/c/cc0461en

2. Verdegem M, Buschmann AH, Latt UW, Dalsgaard AJ, Lovatelli A. The contribution of aquaculture systems to global aquaculture production. Journal of the World Aquaculture Society. 2023 Apr;54(2):206–50. Available from: 10.1111/jwas.12963

3. Debnath PP, Delamare-Deboutteville J, Jansen MD, Phiwsaiya K, Dalia A, Hasan MA, Senapin S, Mohan CV, Dong HT, Rodkhum C. Two-year surveillance of tilapia lake virus (TiLV) reveals its wide circulation in tilapia farms and hatcheries from multiple districts of Bangladesh. Journal of Fish Diseases. 2020 Nov;43(11):1381–9. Available from: 10.1111/jfd.13235

4. Abdelrahman HA, Hemstreet WG, Roy LA, Hanson TR, Beck BH, Kelly AM. Epidemiology and economic impact of disease-related losses on commercial catfish farms: A seven-year case study from Alabama, USA. Aquaculture. 2023 Mar 15;566:739206. Available from: 10.1016/j.aquaculture.2022.739206

5. Dong, H.T., Nguyen, V.V., Le, H.D., Sangsuriya, P., Jitrakorn, S., Saksmerprome, V., Senapin, S. and Rodkhum, C., 2015. Naturally concurrent infections of bacterial and viral pathogens in disease outbreaks in cultured Nile tilapia (Oreochromis niloticus) farms. Aquaculture, 448, pp.427–435. Available from: 10.1016/j.aquaculture.2015.06.027

6. Haenen OL, Dong HT, Hoai TD, Crumlish M, Karunasagar I, Barkham T, Chen SL, Zadoks R, Kiermeier A, Wang B, Gamarro EG. Bacterial diseases of tilapia, their zoonotic potential and risk of antimicrobial resistance. Reviews in Aquaculture. 2023 Feb;15:154–85. Available from: 10.1111/raq.12743

7. Clavijo AM, Conroy G, Conroy DA, Santander J, Aponte F. First report of Edwardsiella tarda from tilapias in Venezuela. Bulletin-European Association of Fish Pathologists. 2002 Jan 1;22(4):280–2.

8. Leal CA, Tavares GC, Figueiredo HC. Outbreaks and genetic diversity of Francisella noatunensis subsp orientalis isolated from farm-raised Nile tilapia (Oreochromis niloticus) in Brazil. Genetics and Molecular Research. 2014 Jul 25;13(3):5704–12. Available from: 10.4238/2014.July.25.26

9. Dong HT, Techatanakitarnan C, Jindakittikul P, Thaiprayoon A, Taengphu S, Charoensapsri W, Khunrae P, Rattanarojpong T, Senapin S. Aeromonas jandaei and Aeromonas veronii caused disease and mortality in Nile tilapia, Oreochromis niloticus (L.). Journal of Fish Diseases. 2017 Oct;40(10):1395–403. Available from: 10.1111/jfd.12617

10. Abu-Elala NM, Abd-Elsalam RM, Younis NA. Streptococcosis, Lactococcosis and Enterococcosis are potential threats facing cultured Nile tilapia (Oreochomis niloticus) production. Aquaculture research. 2020 Oct;51(10):4183–95. Available from: 10.1111/are.147601.

11. Bwalya P, Simukoko C, Hang’ombe BM, Støre SC, Støre P, Gamil AA, Evensen Ø, Mutoloki S. Characterization of streptococcus-like bacteria from diseased Oreochromis niloticus farmed on Lake Kariba in Zambia. Aquaculture. 2020 Jun 30;523:735185. Available from: 10.1016/j.aquaculture.2020.735185

12. Ciji A, Akhtar MS. Stress management in aquaculture: A review of dietary interventions. Reviews in Aquaculture. 2021 Sep;13(4):2190–247. Available from: 10.1111/raq.12565

13. Hasimuna OJ, Maulu S, Nawanzi K, Lundu B, Mphande J, Phiri CJ, Kikamba E, Siankwilimba E, Siavwapa S, Chibesa M. Integrated agriculture-aquaculture as an alternative to improving small-scale fish production in Zambia. Frontiers in Sustainable Food Systems. 2023 May 10;7:1161121. Available from: 10.3389/fsufs.2023.1161121

14. Zhang L, Maulu S, Hua F, Chama MK, Xu P. Aquaculture in Zambia: the current status, challenges, opportunities and adaptable lessons learnt from China. Fishes. 2023 Dec 29;9(1):14. Available from: 10.3390/fishes9010014

15. Oladapo O, Yinusa, M, Bangwe L, Marttin, F, Chimatiro S, Jere N. Zambia - Aquaculture Enterprise Development Project [Internet]. African Development Bank; 2016 [cited 2024 Nov 23]. Available from: https://www.afdb.org/en/documents/document/zambia-aquaculture-enterprise-development-project-93700

16. Siamujompa M, Ndashe K, Zulu FC, Chitala C, Songe MM, Changula K, Moonga L, Kabwali ES, Reichley S, Hang’ombe BM. An investigation of bacterial pathogens associated with diseased Nile tilapia in small-scale cage culture farms on Lake Kariba, Siavonga, Zambia. Fishes. 2023 Sep 8;8(9):452. Available from: 10.3390/fishes8090452

17. Barcellos LG, Nicolaiewsky S, De Souza SG, Lulhier F. The effects of stocking density and social interaction on acute stress response in Nile tilapia Oreochromis niloticus (L.) fingerlings. Aquaculture Research. 1999 Nov;30(11-12):887–92. Available from: 10.1046/j.1365-2109.1999.00419.x

18. Raman RP, Prakash C, Makesh M, Pawar NA. Environmental stress mediated diseases of fish: an overview. Advances in Fish Research. 2013;5:141–58.

19. Team Data. Skalenniveau. DATAtab; 2022.

20. Aboyadak IM, Ali NG, Goda AM, Aboelgalagel WH, Salam A. Molecular detection of Aeromonas hydrophila as the main cause of outbreak in tilapia farms in Egypt. J Aquac Mar Biol. 2015;2(6):2–5.

21. Laltlanmawia C, Ghosh L, Saha RK, Parhi J, Pal P, Dhar B, Saha H. Isolation, identification and pathogenicity study of emerging multi-drug resistant fish pathogen Acinetobacter pittii from diseased rohu (Labeo rohita) in India. Aquaculture Reports. 2023 Aug 1;31:101629. Available from: 10.1016/j.aqrep.2023.101629

22. Sakala T, Mdegela RH, Hangombe BM. Identifying bacteria associated with diseased Oreochromis niloticus in Lake Kariba, Zambia. Available from: https://smujo.id/bbs/article/view/122201.

23. Vaneci-Silva D, Assane IM, de Oliveira Alves L, Gomes FC, Moro EB, Kotzent S, Pitondo-Silva A, Pilarski F. Klebsiella pneumoniae causing mass mortality in juvenile Nile tilapia in Brazil: Isolation, characterization, pathogenicity and phylogenetic relationship with other environmental and pathogenic strains from livestock and human sources. Aquaculture. 2022 Jan 15;546:737376. Available from: 10.1016/j.aquaculture.2021.737376

24. Assane IM, de Sousa EL, Valladão GM, Tamashiro GD, Criscoulo-Urbinati E, Hashimoto DT, Pilarski F. Phenotypic and genotypic characterization of Aeromonas jandaei involved in mass mortalities of cultured Nile tilapia, Oreochromis niloticus (L.) in Brazil. Aquaculture. 2021 Aug 30;541:736848. Available from: 10.1016/j.aquaculture.2021.736848

25. Azzam-Sayuti M, Ina-Salwany MY, Zamri-Saad M, Annas S, Yusof MT, Monir MS, Mohamad A, Muhamad-Sofie MH, Lee JY, Chin YK, Amir-Danial Z. Comparative pathogenicity of Aeromonas spp. in cultured red hybrid tilapia (Oreochromis niloticus× O. mossambicus). Biology. 2021 Nov 17;10(11):1192. Available from: 10.3390/biology10111192

26. Rao S, Pham TH, Poudyal S, Cheng LW, Nazareth SC, Wang PC, Chen SC. First report on genetic characterization, cell-surface properties and pathogenicity of Lactococcus garvieae, emerging pathogen isolated from cage-cultured Cobia (Rachycentron canadum). Transboundary and Emerging Diseases. 2022 May;69(3):1197–211. Available from: 10.1111/tbed.14083

27. Abdel-Latif HM, Khafaga AF. Natural co-infection of cultured Nile tilapia Oreochromis niloticus with Aeromonas hydrophila and Gyrodactylus cichlidarum experiencing high mortality during summer. Aquaculture Research. 2020 May;51(5):1880–92. Available from: 10.1111/are.14538

28. Nicholson P, Mon-On N, Jaemwimol P, Tattiyapong P, Surachetpong W. Coinfection of tilapia lake virus and Aeromonas hydrophila synergistically increased mortality and worsened the disease severity in tilapia (Oreochromis spp.). Aquaculture. 2020 Apr 15;520:734746. Available from: 10.1016/j.aquaculture.2019.734746

